# Whole-genome analysis of multiple wood ant population pairs supports similar speciation histories, but different degrees of gene flow, across their European range

**DOI:** 10.1101/2021.03.10.434741

**Authors:** Beatriz Portinha, Amaury Avril, Christian Bernasconi, Heikki Helanterä, Josie Monaghan, Bernhard Seifert, Vitor C. Sousa, Jonna Kulmuni, Pierre Nouhaud

## Abstract

The application of demographic history modeling and inference to the study of divergence between species is becoming a cornerstone of speciation genomics. The demographic history is usually reconstructed by analysing a single population from each species, assuming that the divergence history inferred between these populations represents the actual speciation history. However, this assumption is rarely explicitly tested, and it may not be met when species diverge with gene flow. For instance, secondary contact between two species after a range expansion may be confined into a specific geographic region. In this study, we tested to what extent the divergence history inferred from two heterospecific populations would vary depending on their geographic locations, using mound-building red wood ants. The wood ant species *Formica polyctena* and *F. aquilonia* have contrasting distributions in Europe and naturally hybridize in Finland. We first performed whole-genome resequencing of 20 individuals sampled in multiple populations across both species ranges. We then reconstructed the divergence histories of distinct heterospecific population pairs using a coalescent-based approach. We found that the analysis of these different population pairs always supported a scenario of divergence with gene flow, suggesting that species divergence started in the Pleistocene (ca. 500 kya) and occurred with continuous asymmetrical gene flow from *F. aquilonia* to *F. polyctena* until a recent time, when migration stopped (2-19 kya, depending on the population pair considered). However, we found support for contemporary gene flow in the sympatric population pair from Finland, where hybrids have been described. Overall, our results suggest that divergence histories reconstructed from a few individuals may be reliable and applicable at the species level. Nonetheless, the geographical context of populations chosen to represent their species should be taken into account, as it may affect estimates of migration rates between species when gene flow is heterogeneous across their geographical ranges.

## INTRODUCTION

Reconstructing divergence histories using genetic data has become gold standard in speciation genomics (Ravinet *et al.*, 2017), which has been eased by the development of sequencing technologies and inference tools (Beichman *et al.*, 2018). Classically, the divergence history between two related species is inferred using samples from a single pair of populations, one from each species, with the purpose of estimating key evolutionary parameters such as divergence times, migration rates, and effective population sizes (e.g., Nadachowska-Brzyska *et al.* 2013; Yagi *et al.*, 2019; Sutra *et al.*, 2019). Such an approach implicitly assumes that the divergence history between the two sampled populations is representative of the divergence history of the species as a whole, i.e. across their whole ranges. This assumption is expected to hold if species diverge in allopatry without gene flow. However, this assumption is rarely explicitly tested, and when divergence occurs with gene flow it is unclear to what extent parameter estimates may fluctuate across both species ranges.

Gene flow between two diverging lineages can vary through time (Sousa & Hey 2013) and space. For instance, secondary contact between two species after a range expansion is likely to only affect populations in a specific geographic region (e.g., Green *et al.*, 2010). In such cases, reconstructing the divergence history between the two species would depend on the geographic location of the set of populations sampled, because populations also evolve in space (see Bradburd & Ralph, 2019 for a recent review on spatial population genetics). While some studies have previously reconstructed the divergence history between several species using multiple population pairs (e.g., Zieliński *et al.*, 2016; Filatov *et al.*, 2016; Pabijan *et al.*, 2017; Rougemont and Bernatchez, 2018; Ito *et al.*, 2020; Garcia-Erill *et al.*, 2021), to our knowledge variation inferred between the multiple comparisons has not been reported. In this study, we investigate how the divergence history between two species may vary depending on the geographic location of the pair of heterospecific populations considered, using mound-building red wood ants.

Red wood ants of the *Formica rufa* group (Hymenoptera, Formicidae) play important ecosystemic roles in boreal forests (Frouz *et al.* 2016; Stockan *et al.* 2016) and are a good system to study variability in the divergence history across species geographical ranges. This is because the Palearctic *F. rufa* group encompasses up to 13 species (Seifert 2021), many with different distribution areas and that likely experienced gene flow or secondary contact in different regions. Phylogenetic studies using mitochondrial markers suggest that speciation within this group took place in the Pleistocene, in the last 500,000 years (Goropashnaya *et al.*, 2004; Goropashnaya *et al.*, 2012). While their speciation history is unknown, they may have diverged in different forest refugia during Pleistocene glaciations (Goropashnaya *et al.*, 2004). Among the *F. rufa* group, *F. polyctena* and *F. aquilonia* are two non-sister species with contrasting distributions. Within the European part of their Palaearctic ranges, *F. aquilonia* shows a boreal-subarctic horizontal and highly montane-subalpine vertical distribution, whereas *F. polyctena* is largely temperate-subboreal and planar to submontane (Seifert, 2018). Both species overlap in Switzerland (where they occupy different altitudinal niches) and around the Baltic sea, and natural hybrids have been characterized in Southern Finland (Kulmuni *et al.*, 2010; Beresford *et al.*, 2017).

Here, we reconstructed the speciation history between *F. polyctena* and *F. aquilonia* using whole-genome data by contrasting multiple pairs of populations sampled from both species across Europe. Our aim was to understand to what extent different population pairs yield similar demographic histories and parameter estimates. The chosen pairs of populations represent situations of present-day strict allopatry (West Switzerland *F. polyctena* vs. Scotland *F. aquilonia* and East Switzerland *F. polyctena* vs. Scotland *F. aquilonia),* allopatry/parapatry in Switzerland (where species are distributed over different altitudinal ranges, but opportunities for gene flow cannot be ruled out completely; Cherix *et al.*, 2012) and sympatry in Finland (where hybridization has been characterized). Our results suggest that divergence started in the Pleistocene and that it occured with continuous asymmetric gene flow from *F. aquilonia* to *F. polyctena.* Interestingly, all population comparisons supported the same divergence scenario, with comparable divergence times ranging from 523,900 to 561,745 years ago, in agreement with previous findings. Nevertheless, we also found differences between population pairs, particularly in contemporary gene flow estimates. Among these differences, we found support for increased bidirectional migration at recent times only in Finland, where gene flow could be mediated by hybrids. By using multiple populations distributed across both species ranges, we were able to both draw a consistent picture of the speciation history (i.e., divergence times, ancestral effective sizes and ancestral migration rates) while uncovering variability in migration rates which could be explained by local opportunities for gene flow.

## MATERIALS AND METHODS

### Study system

*Formica polyctena* and *F. aquilonia* are two Holarctic ant species of the *Formica* genus that inhabit coniferous and mixed forests. They are haplodiploid (females are diploid and males are haploid) and arrhenotokous (mothers produce male offspring from unfertilized eggs; De La Filia *et al.*, 2015). They are social insects and, as such, labour is divided between reproductive queens and workers. As both of these species are polygynous, each nest may have hundreds of egg-laying queens of different ages. The species are also supercolonial, meaning that populations are formed by the association of many cooperating nests (hereafter, population is used as a synonym for supercolony). Polygyny results in low relatedness among individuals sampled within the same nest and/or supercolony (e.g., Sundström *et al.*, 2005). New queens are produced in the Spring and they may shed their wings without a nuptial flight. As such, matings can happen with males from their own population. Finally, if long-range dispersal occurs, it is likely male-biased (Maeder *et al.*, 2016).

### Sampling

Our main aim was to understand to what extent samples from different geographical locations yield similar demographic histories and parameter estimates. To address this, females (workers) of each parental species were sampled from several locations across Europe (Fig. 1). For *Formica polyctena,* sampling was carried out in two locations in Switzerland (East and West) and in the Åland islands (Finland). For each sampling site, three female workers were sampled from different populations, and/or different nests within the same population, whenever possible (Table 1). In addition, one more female of each species was collected in Southern Finland, where hybridization between both species has been previously documented (Beresford *et al.*, 2017). Overall, ten females were sampled for each species (20 individuals in total, Table 1).

**Figure 1.**
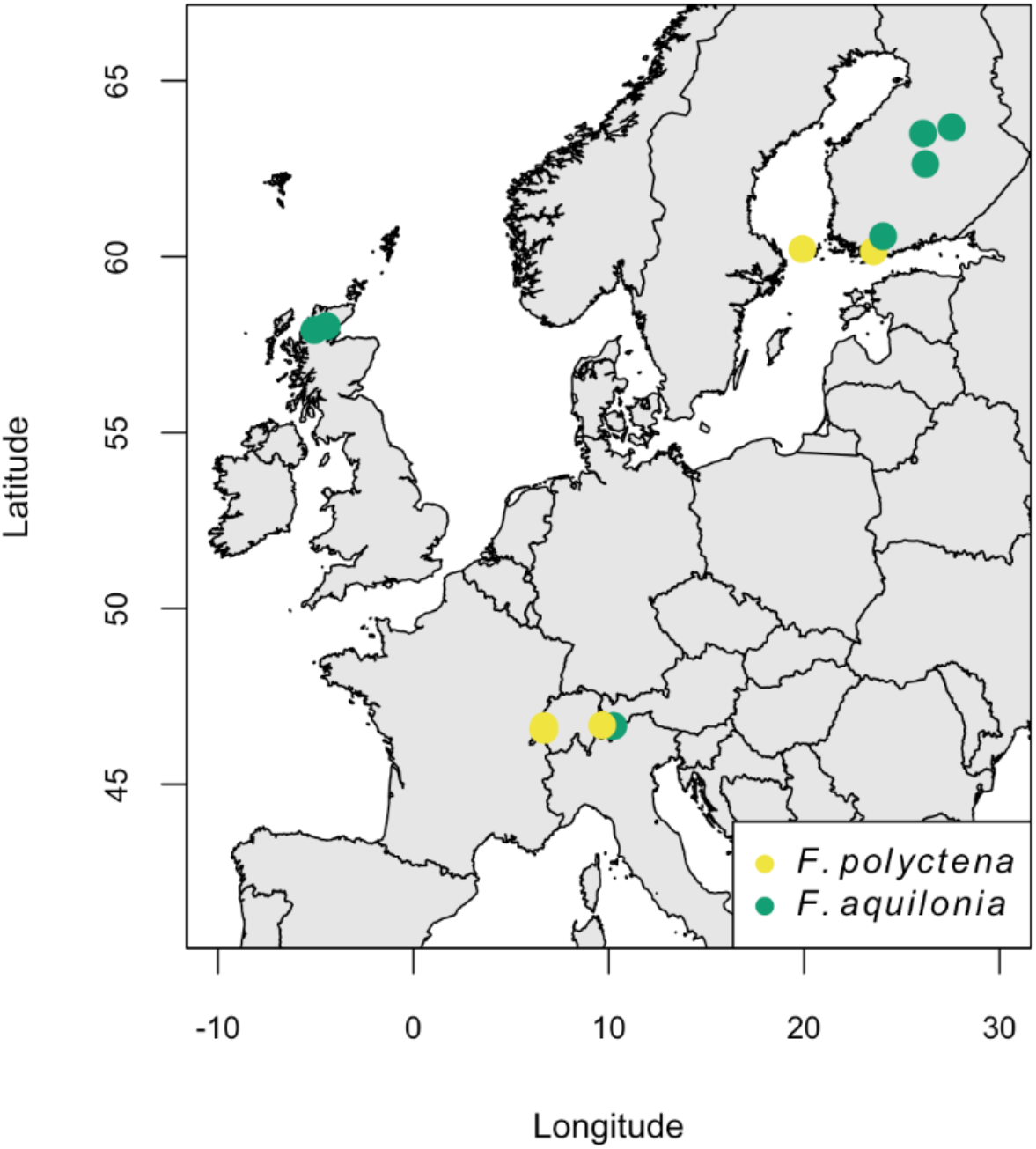
Map of sampling locations. Each symbol represents a sampled individual (some are overlapping).

**Table 1.**
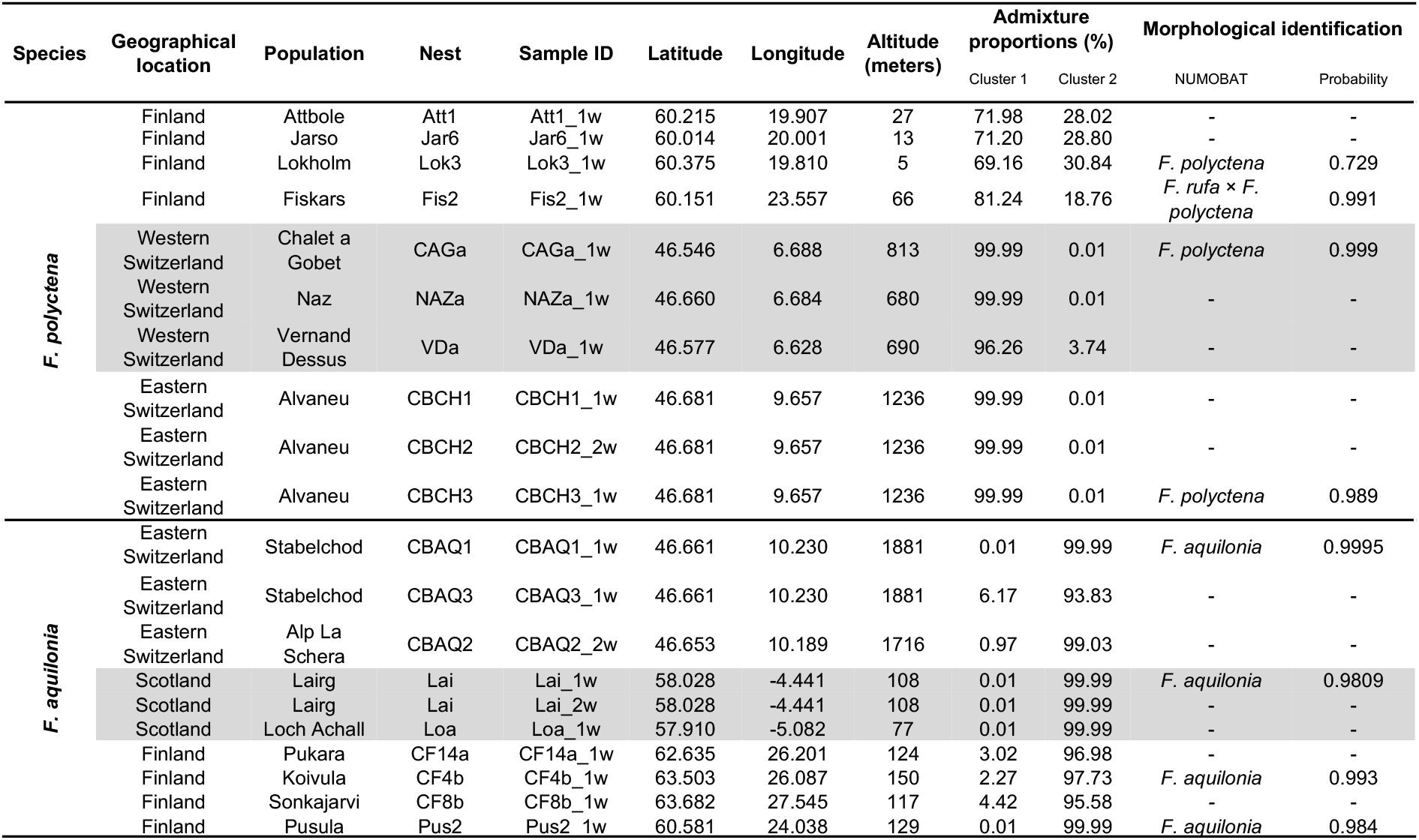
Sample information. Sampling location (locality, geographic coordinates and altitude), nest of origin, ancestry proportions (reconstructed by the sNMF analysis for K=2), and assignment probabilities for the 20 sampled individuals.

### Morphological identification

Prior to sequencing, a subset of samples (at least one nest per geographical location, Table 1) were morphologically identified at the species level using Numeric Morphology-Based Alpha-Taxonomy (NUMOBAT). Overall, 16 morphological characters were investigated (CS, CL/CW, SL/CS, nCH, OccHL, nGU, GUHL, nPN, mPnHL, nMes, nMet, MetHL, nPr, EyeHL, nSc and SL/Smax, Seifert, 2018).

### DNA extraction and sequencing

For all samples, DNA was extracted from whole-bodies with a SDS (sodium dodecyl sulfate) protocol. DNA libraries were constructed using NEBNext DNA Library Prep Kits (New England Biolabs). Samples were processed and sequenced at Novogene (Hong Kong) as part of the Global Ant Genomics Alliance (Boomsma *et al.*, 2017). Whole-genome sequencing was performed on Illumina Novaseq 6000 (150 base pairs paired-end reads), targeting 15× per individual.

Raw Illumina reads and adapter sequences were trimmed using Trimmomatic (v0.38; parameters LEADING:3, TRAILING:3, MINLEN:36; Bolger *et al.*, 2014) before mapping against the reference genome (Nouhaud *et al.*, 2021) using BWA MEM with default parameters (v0.7.17; Li and Durbin, 2010). Duplicates were removed using Picard Tools with default parameters (v2.21.4; http://broadinstitute.github.io/picard).

### SNP calling and filtering

Single nucleotide polymorphisms (SNPs) and genotypes were called across all samples with freebayes (v1.3.1; Garrison and Marth, 2012), disabling population priors (-k). After SNP calling, the VCF was normalised using vt (v0.5; Tan *et al.*, 2015) and sites located at less than two base pairs from indels were excluded, along with sites supported by only Forward or Reverse reads. Multi-nucleotide variants were decomposed with the vcfallelicprimitives command from vcflib (v1.0.1).

Only biallelic SNPs with quality equal or higher than 30 were kept. In order to refrain from removing entire sites when only a subset of individuals had inadequate genotype calls, individual genotypes with genotype qualities lower than 30 were coded as missing data. Genotypes with depth of coverage lower than eight were also coded as missing data, after which sites with missing data across more than half of our samples were removed.

To remove genotyping errors that cause sites to show excessive heterozygosity (e.g., due to unresolved paralogs or alignment errors), we first pooled all samples together, creating a Wahlund effect on purpose, after which we applied a filter based on Hardy-Weinberg Equilibrium and excluded sites displaying heterozygote excess (*p* < 0.01; see e.g., Pfeifer *et al.*, 2018).

We applied a filter based on sequencing depth by setting individual-specific thresholds: sites were only kept if their coverage was between half and twice the mean individual value. Finally, we removed sites located on the third chromosome, also known as the social chromosome. This chromosome harbours genes responsible for polymorphism in social organization in *Formica* species, controlling if a colony is headed by one (monogynous) or multiple (polygynous) queens (Brelsford *et al.*, 2020). Recombination is rare between monogynous and polygynous alleles of this chromosome (supergene, Brelsford *et al.*, 2020), leading to the maintenance of ancestral polymorphisms across *Formica* species which could bias our demographic inference.

### Population structure

Population structure was studied by means of two individual-based methods, Principal Component Analysis (PCA) and sNMF clustering analysis (Frichot *et al.*, 2014), the latter of which estimates individual ancestry coefficients. These analyses were performed in R (v3.6.3; R Core Team, 2017) using the LEA package (v3.0.0; Frichot and François, 2015). The sNMF analysis was repeated 20 times for a number of ancestral clusters (K) ranging from 2 to 8. The K value with lowest cross-entropy obtained by sNMF was considered as the best K value, and the run with the lowest cross-entropy for the best K value was considered as the best run.

Observed and expected heterozygosity, inbreeding coefficients (*F*_IS_) and pairwise fixation indices (*F*_ST_; Weir and Cockerham, 1984) were calculated using custom scripts. Pairwise *F*_ST_ values were also computed between populations using the SNPRelate package (v1.20.1; Zheng *et al.*, 2012).

### Demographic modelling

To document the divergence history across the species ranges, we compared alternative demographic models to demographic parameters inferred from the site frequency spectrum (SFS) obtained from different combinations of populations. This was done using the composite-likelihood method implemented in fastsimcoal2 (v2.6; Excoffier *et al.*, 2013, parameters detailed in Supplementary Table 1). For allopatric cases, we considered three population pairs: *F. polyctena* from West Switzerland vs. *F. aquilonia* from Scotland; *F. polyctena* from East Switzerland vs. *F. aquilonia* from Scotland and *F. polyctena* from West Switzerland vs. *F. aquilonia* from East Switzerland. The sympatric population pair was formed by Finnish populations of both species. Each model was run 100 times, with 80 iterations per run for likelihood maximization, and 200,000 coalescent simulations per iteration to approximate the expected SFS. The mutation rate was assumed as 3.5×10^-9^ per bp per haploid genome per generation, which is approximated from estimates available for social insects (Liu *et al.*, 2017). No population growth was assumed (i.e., population growth rates were zero).

In the *Formica* genus, young queens can start laying eggs in their first years of life and have been estimated to live up to five years for *F. polyctena* (Horstmann, 1983), with queens of different ages within a single nest (i.e., overlapping generations). As such, generation time was assumed to be 2.5 years.

After obtaining point parameter estimates and expected likelihoods for all models (see below) tested with all population comparisons considered, the model with the highest expected likelihood was chosen as the best model in each case.

### Speciation history between *F. polyctena* and *F. aquilonia*

To ascertain whether the speciation history between *F. aquilonia* and *F. polyctena* is different across both species ranges, we considered four divergence scenarios: allopatry, sympatry, isolation after migration, and migration after isolation (Fig. 2). The first two models allow for populations to change sizes at any time (Fig. 2A,B), while the remaining models allow resize events only when migration changes (Fig. 2C,D). The model with a scenario of sympatry between the populations (Fig. 2B) allows the migration rates between populations to change after the population sizes change. Parameter search ranges were improved for this particular model based on preliminary analyses, in order to decrease the probability that the parameter estimates obtained represent local maxima.

**Figure 2.**
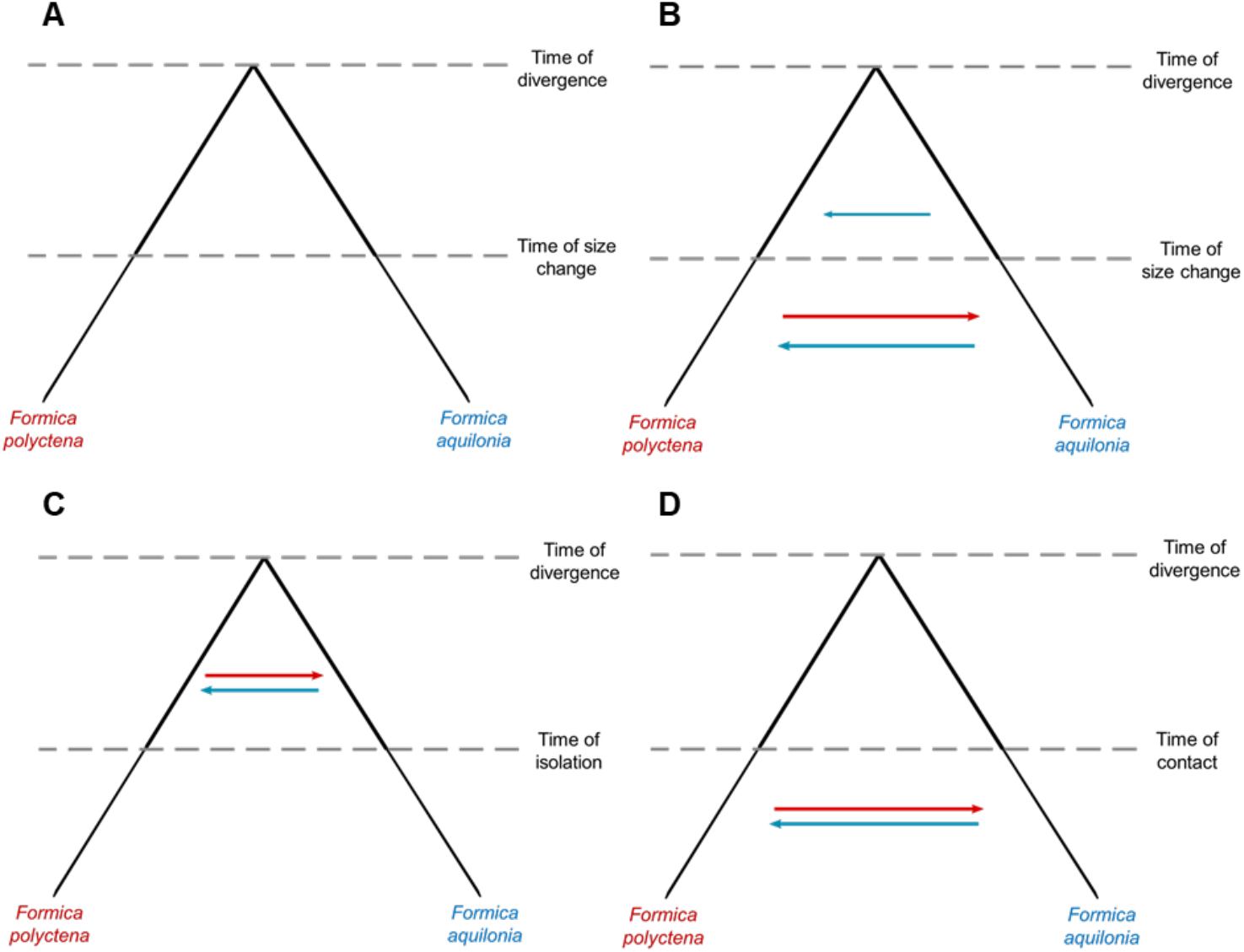
Demographic models designed to study the speciation history between *Formica polyctena* and *F. aquilonia.* **A** Allopatry scenario. **B** Sympatry scenario, allowing for different migration rates throughout the divergence of the populations. **C** Isolation after migration scenario. **D** Migration after isolation scenario. Arrows represent migration. The direction of gene flow is indicated by the direction and colour of the arrows (red represents gene flow from *F. polyctena* into *F. aquilonia;* blue represents gene flow from *F. aquilonia* into *F. polyctena).* The different thickness in the lines representing the populations represent changes in effective size, which can happen either by instantaneous contractions or expansions.

### Ghost introgression

In order to rule out the possibility that the signal of gene flow we detect between *F. aquilonia* and *F. polyctena* is actually caused by gene flow from an unsampled (so-called ghost) species into either of the focal species, we modeled two different ghost scenarios which are based on species relationships within the *F. rufa* group as described in Goropashnaya *et al.* (2012). The first scenario models ghost introgression from *F. rufa,* which is a sister species to *F. polyctena,* and which may send migrants to either *F. polyctena* (Fig. 3A), or *F. aquilonia* (Fig. 3C). The second, alternative scenario models ghost introgression from either *F. lugubris* and/or *F. paralugubris,* which are in the same clade as *F. aquilonia,* and which may send migrants to either *F. aquilonia* (Fig. 3B), or *F. polyctena* (Fig. 3D). These models purposefully did not consider direct migration between *F. polyctena* and *F. aquilonia,* since they were designed to rule out the possibility that the signal of gene flow between the focal species could be caused by gene flow from an unsampled sister species.

**Figure 3.**
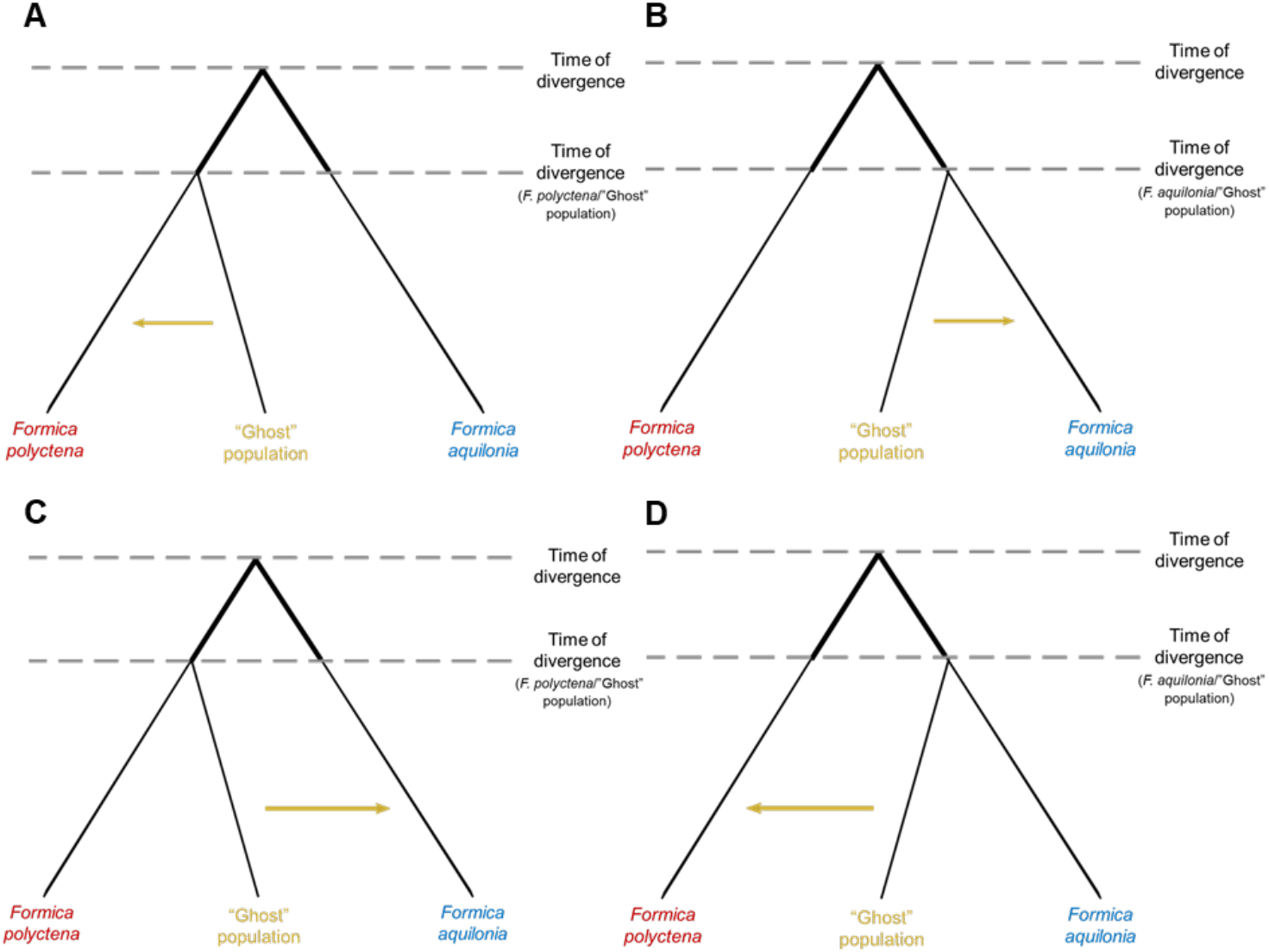
Demographic models designed to study possible introgression from unsampled species (“ghost”) into *Formica polyctena* or *F. aquilonia.* **A** Scenario where the unsampled species is divergent from *F. polyctena,* into which it sends migrants. **B** Scenario where the unsampled species is divergent from *F. aquilonia,* into which it sends migrants. **C** Scenario where the unsampled species is a sister of *F. polyctena* and sends migrants into *F. aquilonia.* **D** Scenario where the unsampled species is a sister of *F. aquilonia* and send migrants into *F. polyctena.* Gene flow and changes in population sizes are depicted as in Figure 2.

### SFS characteristics

To perform the demographic analyses detailed above, we built SFSs using data from two populations (2D-SFS). As we lack a good outgroup to infer the ancestral state of the alleles in our dataset, folded SFSs were built using the minor allele frequency (MAF) method. We ensured that there was no missing data by downsampling genotypes. To do this, a minimum sample size across all sites was determined (corresponding to the number of individuals to resample from minus the maximum number of missing data per site). Individuals were resampled in windows of 50 Kbp, discarding blocks where the mean distance between consecutive SNPs in a given block was less than 2 bp. To maximize the number of sites that could be kept, the individuals selected to be resampled in each window were the ones with higher amounts of data in that specific window.

All SFSs included the number of monomorphic sites, which, in conjunction with a mutation rate, allows scaling parameter estimates inferred by the models (e.g. to obtain divergence times in number of generations). We estimated these numbers using the proportion of polymorphic sites in relation to the total number of callable sites of individuals in a specific dataset. The total number of callable sites was obtained for each individual using mosdepth (v0.2.9; Pedersen & Quinlan, 2017) considering individual depth of coverage thresholds defined earlier for SNP calling.

## RESULTS

### Sequencing and SNP calling

Illumina sequencing yielded on average 7.12 Gb of raw data per sample (min: 5.87, max: 8.29). After trimming, mapping and filtering, average sequencing depth was 15.6× ± 1.66 (standard deviation). SNP calling using Freebayes recovered 2,856,374 biallelic sites with quality values above 30. Among these sites, 2,211,441 were left after filtering on sequencing depth, individual coverage and excessive heterozygosity, and 2,054,352 after removing sites located on the third chromosome. The fraction of missing data per site averaged 15.49% across all 20 individuals in the final SNP set.

### Morphological species identification

A subset of samples used for genomic analyses were also used in morphological species identification. The analysis of 16 morphological characters under the NUMOBAT framework supported our prior species assignment for all *F. aquilonia* samples and non-Finnish *F. polyctena* samples. For these samples, all posterior probabilities of the morphological assignment were greater than 98% (Table 1). Individual samples collected in Åland (Lokholm) were morphologically assigned as *F. polyctena* (72.9%) and *F. polyctena* × *F. aquilonia* hybrids (26.8%), while samples collected in Fiskars were assigned as *F. polyctena* × *F. rufa* hybrids (99.1%, Table 1). The clustering analysis based on the genome-wide data carried with sNMF assigned these two individuals (Lok3_1w and Fis2_1w, respectively; Table 1) as admixed *F. polyctena* (69.16 and 81.24% assignment proportions, respectively; see below and Table 1). As such they were included in all analyses with the prior that they may be admixed themselves.

### Summary statistics and genetic structure

Genome-wide, average pairwise differentiation indices (*F*_ST_) for all possible combinations of geographic sampling locations were moderately high (*F*_ST_ > 0.1) in all cases (Table 2). The highest value was recorded between *F. polyctena* individuals from East Switzerland and *F. aquilonia* individuals from Scotland (*F*_ST_=0.497), and the lowest between the *F. polyctena* individuals from Finland and West Switzerland (*F*_ST_=0.113). Average differentiation between intraspecific samples of the parental species was 0.202 for *F. aquilonia* and 0.131 for *F. polyctena.* Interspecific differentiation ranged from 0.256 to 0.497. Average interspecific differentiation was 0.398.

**Table 2.**
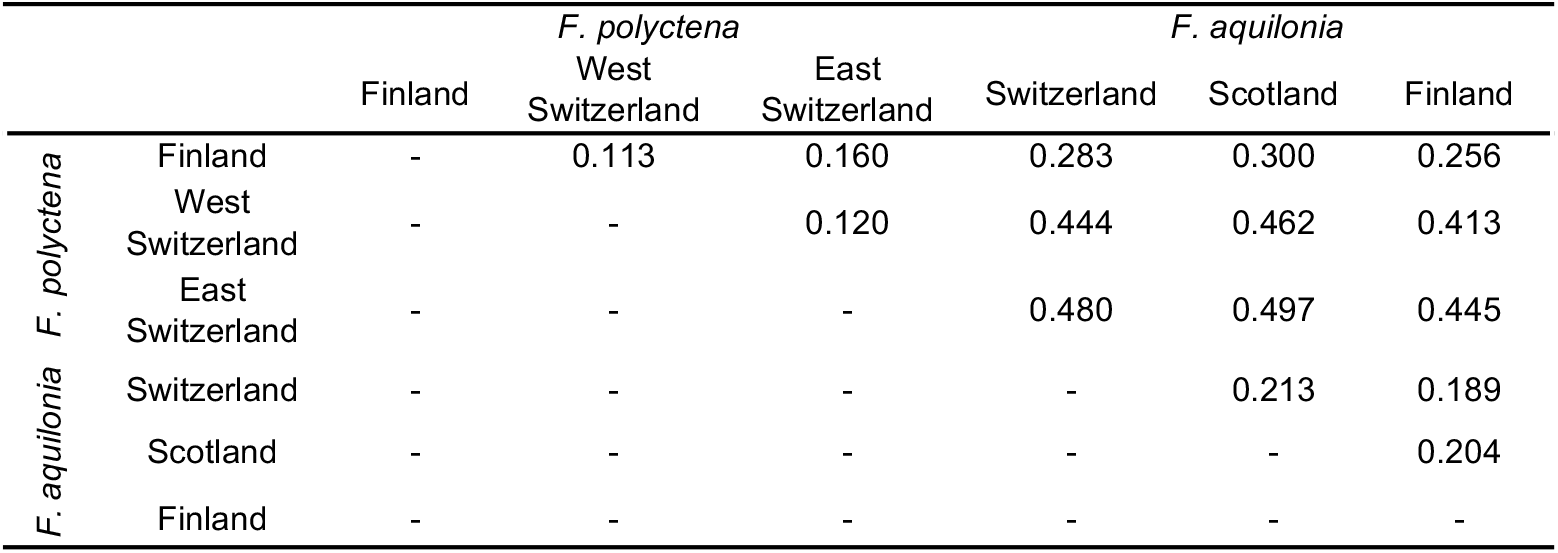
Pairwise fixation indexes (*F*_ST_) between geographic sampling locations of *Formica polyctena* and *F. aquilonia* used in this study.

PCA clearly separated the parental species into two ends of the first principal component (PC, Fig 4). PC1 explained ~29% of the variation in the data and was statistically significant (*p* < 0.01; Supplementary Figure 1). Finnish individuals of *F. polyctena* were plotted closer to *F. aquilonia* individuals, when compared to other non-Finnish *F. polyctena* individuals.

**Figure 4.**
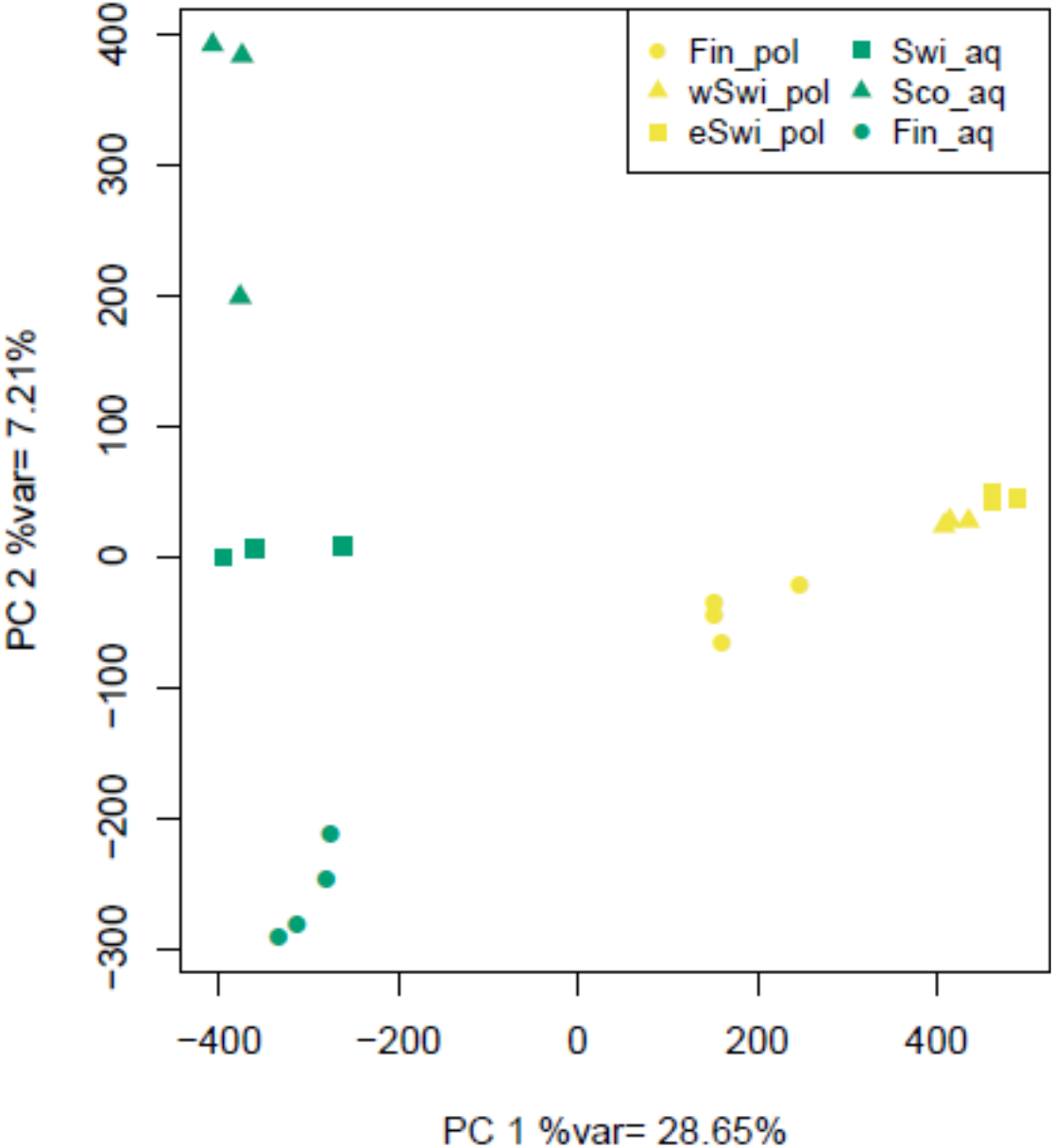
Principal Component Analysis. Principal Component (PC) 1 is shown plotted against PC2. *Formica polyctena* individuals are shown in yellow, with *F. aquilonia* females shown in green. Each point represents an individual and the different point symbols represent the different geographic sampling locations. Abbreviations are as follows: Fin_pol = *F. polyctena* in Finland; wSwi_pol = *F. polyctena* in West Switzerland; eSwi_pol = *F. polyctena* in East Switzerland; Swi_aq = *F. aquilonia* in Switzerland; Sco_aq = *F. aquilonia* in Scotland; Fin_aq = *F. aquilonia* in Switzerland

The sNMF analysis considered two to eight possible ancestral clusters (K). Cross entropy analysis (Supplementary Figure 2) revealed that the best K value for our data was two (Table 1, Supplementary Figure 3). In this case, individuals of both parental species clustered with each other. Interestingly, Finnish individuals of each parental species show some ancestry from the opposite ancestral cluster, up to an ancestry proportion of ~31% from the *F. aquilonia* ancestral cluster in Finnish *F. polyctena* and ~4.42% from the *F. polyctena* ancestral cluster in Finnish *F. aquilonia.* Overall, the sNMF analysis agreed with the morphological identification carried with 16 characters.

Mean expected heterozygosity (*H*_e_) per sampling location ranged from 0.120 to 0.185 (Table 3). *F. aquilonia* from Scotland had the lowest value (0.120), while *F. polyctena* from Finland had the highest *H*_e_ (0.185). Mean observed heterozygosity (*H*_o_; Table 3) per location ranged from 0.103 (Scotland *F. aquilonia*) to 0.169 (Finland *F. polyctena*). All populations, excluding *F. polyctena* from East Switzerland, had lower observed than expected heterozygosity values, translating to positive inbreeding coefficients (*F*_IS_, Table 3). The heterozygosity values are not strictly comparable across locations, since some locations have all samples coming from different populations, whereas other locations consist of samples from the same population (Table 1).

**Table 3.**
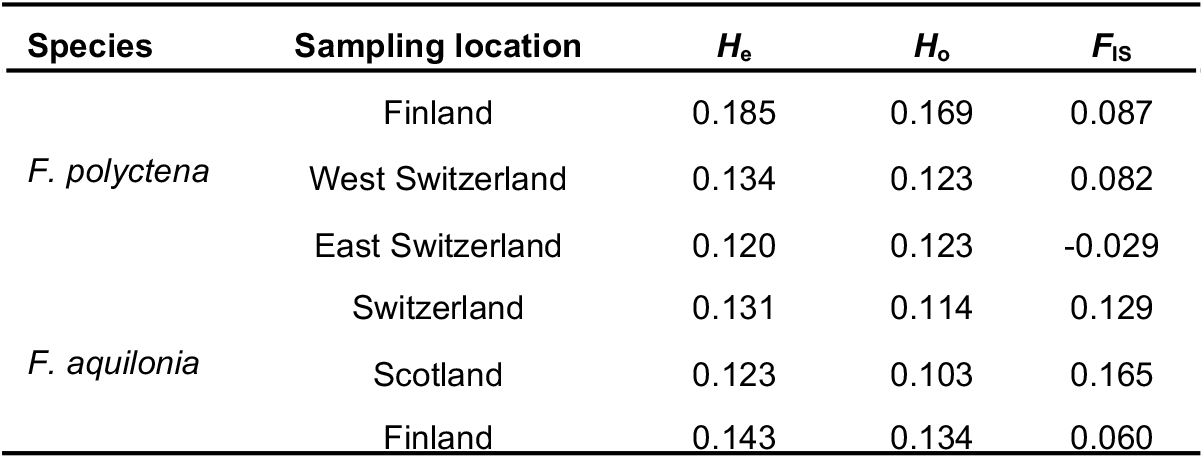
Mean expected (H_e_) and observed (H_o_) heterozygosities and mean inbreeding coefficient (F_IS_) of the geographic sampling locations.

### All comparisons support a scenario of divergence with gene flow

In order to study the speciation history between *F. polyctena* and *F. aquilonia,* we first analysed locations where no hybridization had been previously detected, focusing on the following pairs: (i) *F. polyctena* from Western Switzerland vs. *F. aquilonia* from Scotland (allopatric), (ii) *F. polyctena* from Eastern Switzerland vs. *F. aquilonia* from Scotland (allopatric) and (iii) *F. polyctena* from Western Switzerland vs. *F. aquilonia* from Eastern Switzerland (allopatric/parapatric). For all these population comparisons, the model that implemented divergence in sympatry was found to be the best fit (Fig. 6; Supplementary Tables 2-4 for parameter estimates obtained with all models). The direction of gene flow, migration rates, divergence times and ancestral population sizes were consistent between comparisons involving different pairs of locations (Fig. 5, panels A, B and C).

**Figure 5.**
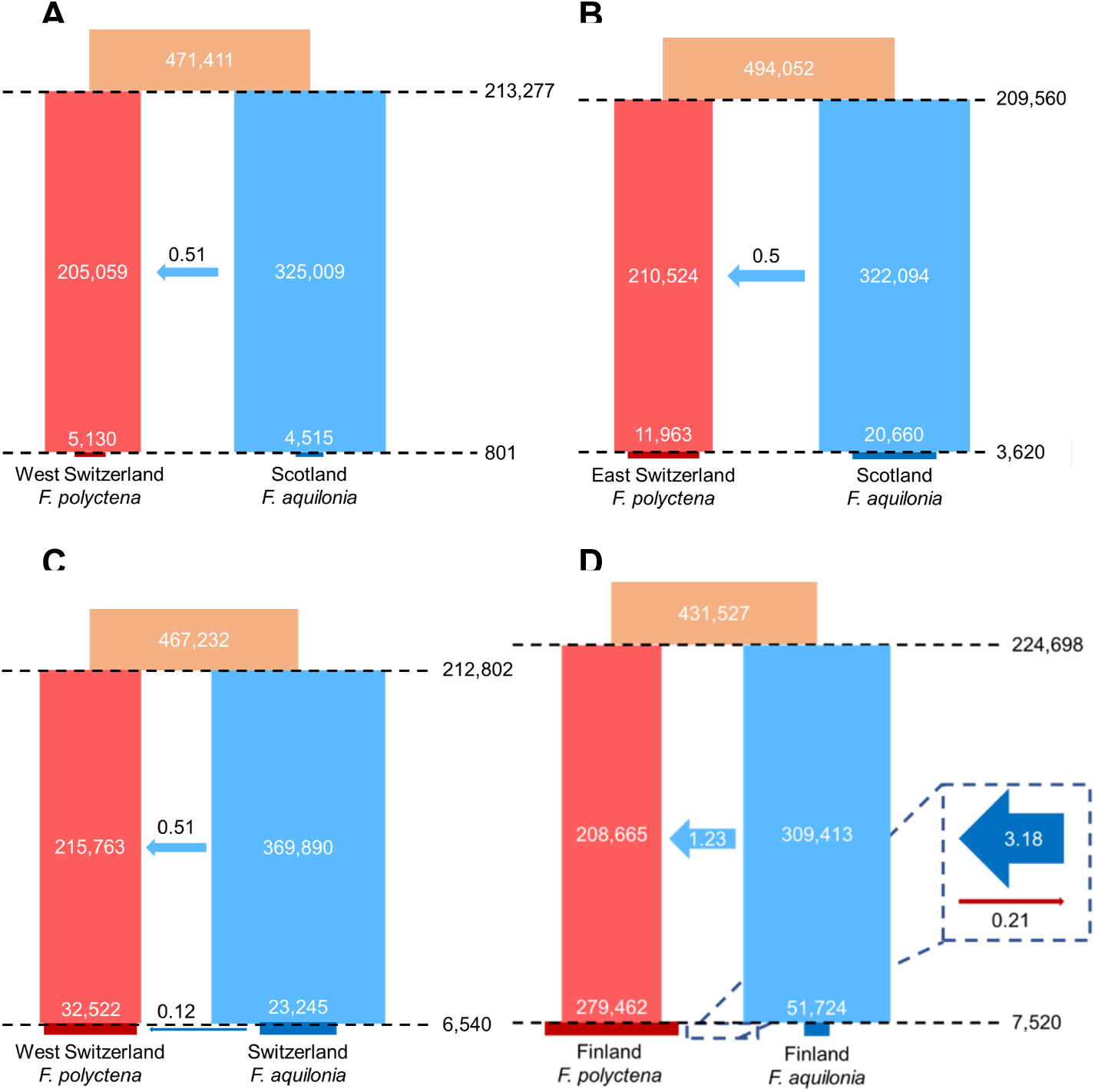
Multiple sample pairs suggest a similar speciation history between *Formica polyctena* and *F. aquilonia.* Best divergence scenarios are depicted for the West Switzerland *F. polyctena* vs. Scotland *F. aquilonia* comparison (**A**), the East Switzerland *F. polyctena* vs. Scotland *F. aquilonia* comparison (**B**), the West Switzerland *F. polyctena* vs. Switzerland *F. aquilonia* comparison (**C**) and the Finland *F. polyctena* vs. Finland *F. aquilonia* comparison (**D**). All times are given in number of generations and represented proportionally to each other across panels, as the time of divergence in panel A was taken as reference. All effective sizes are given in number of haploids. Sizes at a given time are represented proportionally to each other across panels, with the *F. polyctena* sizes in panel A serving as reference (i.e., all recent sizes are proportional to each other but not to ancestral sizes, while all ancestral sizes are proportional to each other but not to recent sizes). Arrows indicate the number of migrants per generation, their size is representative of this value. The direction and colour of the arrows are indicative of the direction of the gene flow.

First, gene flow between species was inferred to be asymmetrical, with migrants moving exclusively from *F. aquilonia* into *F. polyctena* at a rate averaging 0.5 migrants per generation *(2Nm*). Second, the time at which the populations of each species diverged was consistent across the different comparisons and ranged between 209,560 generations (Eastern Switzerland *F. polyctena* vs. Scotland *F. aquilonia*) and 213,277 generations (Western Switzerland *F. polyctena* vs. Scotland *F. aquilonia*). Assuming a generation time of 2.5 years, the estimates for the divergence between these species ranged from 523,900 to 533,193 years ago. Third, the effective size of the ancestral population of both species was consistently estimated to be between 460,000 and 500,000 haploid individuals across comparisons. After divergence, *F. aquilonia* was consistently inferred to have a larger *N*_e_ than *F. polyctena* throughout their history. Finally, our results for all comparisons indicate that both species undergo simultaneous population size contractions which would have occurred between 801 (Western Switzerland *F. polyctena* vs. Scotland *F. aquilonia*) and 6,540 generations ago (Western Switzerland *F. polyctena* vs. Eastern Switzerland *F. aquilonia*). In years, these estimates place the times of population decline between 2,003 and 16,350 years ago.

### Sympatric Finnish samples support a similar speciation history, but with recent bidirectional gene flow

As *F. aquilonia* and *F. polyctena* hybridize in Finland, we tested whether this could have an impact on the divergence history inferred when analyzing a pair of populations from this area. While the expected likelihood of the migration after isolation model was the highest for this population comparison (Supplementary Table 5 for parameter estimates obtained with all models), it was only 3.65 log units better than the likelihood of the model that considers divergence in sympatry. This suggests that both models fit this dataset equally well. As the parameter estimates (e.g., times of divergence and size change, ancestral population sizes) obtained under the sympatry scenario are comparable to those obtained with the comparisons outside of Finland, we consider that the sympatry model is a more parsimonious description of the demographic history between the Finnish *F. polyctena* and *F. aquilonia* individuals (Fig. 6D). The time of divergence estimated under the sympatry scenario for the Finnish comparison, 224,698 generations (561,745 years), was in line with previous estimates. The size of the ancestral population of both species in the model was also comparable to the estimates obtained for the other comparisons (431,527 haploid individuals), and the ancestral population sizes followed the previous trends, with larger estimates for *F. aquilonia* (309,413 haploids) than *F. polyctena* (208,665). However, the size of the Finnish *F. polyctena* population increased in more recent times, while Finnish *F. aquilonia* still contracted, similar to non-Finnish *F. aquilonia*. The time of size change was estimated at 7,520 generations (18,800 years), slightly older than what was inferred for other comparisons. As observed for non-Finnish samples, prior to the population resizes there was unidirectional gene flow from *F. aquilonia* into *F. polyctena* in Finland. However, the migration rate was inferred at 1.23 migrants per generation, which was greater than the average *2Nm* = 0.5 inferred from non-Finnish comparisons (see above). Moreover, in recent times, bidirectional gene flow was inferred, with 3.18 haploid migrants per generation moving from *F. aquilonia* into *F. polyctena* and 0.21 haploid individuals migrating in the other direction.

### Past gene flow between *F. polyctena* and *F. aquilonia* cannot be explained by migration from an unsampled sister species

As several species from the *F. rufa* group are known to hybridize, we tested whether the patterns of gene flow inferred between *F. polyctena* and *F. aquilonia* could be caused by migration from an unsampled, “ghost” species. The expected likelihood of these models with “ghost” introgression was lower than all other “non-ghost” models, suggesting that gene flow between *F. polyctena* and *F. aquilonia* during divergence was the most parsimonious scenario for the samples considered in our study (Supplementary Tables 2 to 5 for the parameter estimates obtained with both models for all population pairs).

## DISCUSSION

Divergence history between two related species is often inferred using samples from a single population pair assuming these samples capture the divergence history throughout the species’ ranges. Yet, this assumption is rarely tested. In this study, we test this assumption by using whole-genome data from samples collected across the European distributions of *Formica polyctena* and *F. aquilonia* to reconstruct their speciation history. Using a moderate number of individuals from each geographic sampling site, we were able to infer consistent speciation histories across distinct sample pairs, both in terms of mode of divergence and parameter estimates (divergence times, ancestral population sizes). As we compared several different pairs of heterospecific populations, our approach also underlined how present-day species distributions affect the inference of the divergence history. Interestingly, we detected reduced gene flow at recent times between allopatric sampling locations while gene flow increased between sympatric Finnish populations in recent times. Finally, our coalescent simulations ruled out the alternative scenarios where gene flow inferred between *F. polyctena* and *F. aquilonia* would be actually caused by gene flow from a third, unsampled species. Below, we discuss the implications of our findings regarding divergence in the *F. rufa* group, and, more generally, the insights gained from contrasting two species across their ranges in terms of speciation research.

### Samples from multiple locations yield a similar divergence history between *F. polyctena* and *F. aquilonia*

All analyses carried out with several heterospecific samples across the species ranges supported the same scenario: *F. polyctena* and *F. aquilonia* diverged with gene flow. Formerly, the possibility of divergence in allopatry in different glacial refugia had been discussed for *F. rufa* group ants (Goropashnaya *et al.*, 2004). However, it is not surprising that we found evidence for recent gene flow between *F. polyctena* and *F. aquilonia,* as many *Formica* species retain the ability to interbreed and produce viable offspring (Seifert and Goropashnaya, 2004, Kulmuni *et al.* 2010, Purcell *et al.*, 2016) and interspecific gene flow has been previously described in many ant species (Feldhaar *et al.*, 2008; Seifert, 2009; Steiner *et al.*, 2011; Seifert, 2019). Yet, one of our most unexpected results is that gene flow during divergence is consistently inferred to be asymmetrical, with gene flow from *F. aquilonia* to *F. polyctena*. This would be observed if prezygotic isolation mechanisms were stronger in *F. aquilonia* than in *F. polyctena* or if *F. aquilonia* was more likely to disperse than *F. polyctena*. This signal of unidirectional gene flow could also be linked to a difficulty for *F. polyctena* individuals to find conspecific mates, which would be expected if this species has a smaller population size than *F. aquilonia.*

While the analyses of multiple heterospecific sample pairs supported the same divergence scenario, they also yielded similar divergence time estimates. The results obtained across all comparisons dated the divergence time between *F. polyctena* and *F. aquilonia* around 540,000 years on average, placing the divergence between both species in the Pleistocene. Our results are in line with previous estimates obtained using mitochondrial markers which dated the diversification of the *F. rufa* group (including *F. polyctena* and *F. aquilonia*) around 490,000 years ago (Goropashnaya *et al.* 2004, 2012). In all heterospecific population pairs considered, the effective size of the ancestral population of both *F. polyctena* and *F. aquilonia* was inferred between 460,000 and 500,000 haploid individuals. After the divergence of these species, *F. aquilonia* was always inferred to have a larger ancestral *N*_e_ than *F. polyctena* and consistent *N*_e_ estimates were obtained across models, indicating that samples of each species from different locations likely shared the same ancestral population. Due to the supercoloniality of both *F. polyctena* and *F. aquilonia*, we suggest that these species could follow the dynamics of a metapopulation (i.e. different supercolonies would be demes within a metapopulation). This could inflate our effective population size estimates, as coalescence of lineages within the same deme is expected to be faster than between lineages in different demes in a metapopulation (Wakeley, 2004). Under this scenario, lineages would be “trapped” within their demes before being able to coalesce with lineages from distinct demes. The maintenance of these lineages over longer time scales would therefore inflate the estimated effective population sizes. However, as both species in our study have similar colony structures, they should be equally impacted by this overestimation.

While the two study species are known to hybridize in Southern Finland (Kulmuni et al. 2010), they can also hybridize with other closely related species from the *F. rufa* group such as *F. lugubris, F. paralugubris* or *F. rufa* (Seifert and Goropashnaya, 2004; Seifert *et al.*, 2010, Seifert 2021). In our ghost introgression models, we tested all possible scenarios of gene flow from an unsampled *F. rufa* group species into either *F. polyctena* or *F. aquilonia*, considering no direct gene flow between our two focal species. We did this to rule out the possibility that the signal of gene flow between our focal species is caused by gene flow from another, unsampled species. Under this hypothesis, the signal of gene flow from *F. aquilonia* into *F. polyctena* that we observe would be actually caused by gene flow from a sister species to *F. aquilonia* into *F. polyctena.* Our results indicate that such unaccounted migration from an unsampled species could not explain the observed pattern of migration between our two focal species. However, our approach relied on modeling unsampled lineages, where coalescent events might happen. In the future, these results should be confirmed by collecting the remaining species of the *F. rufa* group and reconstructing the speciation history of the group as a whole, allowing for gene flow from multiple lineages into the same focal species.

This work represents to our knowledge the first reconstruction of speciation histories between two ant species using genome-wide data. However, as for many non-model organisms, some key parameters required for inference remain unknown. The generation time we used was based on *F. polyctena* queen longevity estimates (Horstmann, 1983), while mutation rate was approximated using data from other social insects (honeybee and bumblebee mutation rates; reviewed in Liu et al., 2017). While these approximations should not bias our inferences, some uncertainty is associated with the date estimates provided throughout our manuscript.

### Present-day context of heterospecific sample pairs induces variability in inferred migration rates

By comparing samples from multiple locations across both species ranges, our approach pinpointed commonalities in the divergence histories across all comparisons, providing a robust picture of the speciation history between *F. polyctena* and *F. aquilonia*. It also allowed us to document variation in the inferred estimates of certain parameters. As such, we can interpret this observed variation in parameter estimates in light of the present-day context of the heterospecific populations under consideration (i.e., whether or not they are presently in contact).

The Finnish *F. polyctena* population is clearly different where it comes to its recent N_e_. Instead of estimates indicating a contraction at the time of size change, akin to the other populations of both *F. polyctena* and *F. aquilonia,* estimates suggest that this population has recently expanded. This seems unlikely, given that *F. polyctena* in Finland is at its range margins (Stockan & Robinson 2016) and based on a national survey is the minority species in Finland (Punttila & Kilpeläinen 2009). We suggest it is more likely that the signal of population expansion in Finnish *F. polyctena* is actually caused by gene flow.

Admixture between *F. aquilonia* and *F. polyctena* in Southern Finland is also supported by our results regarding population structure. Interspecific differentiation between the Finnish populations of *F. polyctena* and *F. aquilonia* was reduced and ancestry coefficients also suggested moderate introgression between both species in Finland. Finnish samples were also more genetically diverse than their conspecific populations sampled outside Finland. Particularly, the Finnish *F. polyctena* had the highest mean expected heterozygosity of all sampled populations, in line with the larger effective size inferred with our coalescent simulations.

The bidirectional contemporary gene flow detected in Finnish populations could be expected under two non-mutually exclusive scenarios. The first is direct introgression of alleles from *F. polyctena* into *F. aquilonia,* which could be facilitated by human activity. The forest management strategy practiced in Finland results in the formation of sharp boundaries between habitats more suitable for *F. aquilonia* (old forests with shade and humidity) and areas where *F. polyctena* can establish, e.g. forest edges, as it can withstand increased exposure to sunlight (Punttila, 2020). This phenomenon might increase opportunities for direct contact between these two species due to closer proximity between heterospecific nests. The second scenario would involve the *F. polyctena* × *F. aquilonia* hybrid populations, which are common in Southern Finland (Beresford *et al.*, 2017). These hybrid populations could mediate gene flow between the species via backcrosses between hybrids and individuals of the parental species (e.g., indirect gene flow). Elucidating these causes would require a dense survey of both parental and hybrid wood ant populations in Finland.

Variation in recent population effective sizes is to be expected as the current populations of these species would have evolved independently from each other, once the ancestral population of each species split into the ancestors of the sampled populations. Variation in recent *N*e estimates obtained for the same population when compared to different heterospecific populations is likely caused by variation in the estimated time at which the size of the populations changes. When the time of size change is estimated to be older, the post-resize populations are estimated to have larger effective sizes.

### Implications for speciation research

In this study, we used red wood ants to investigate how the divergence history between two species may vary depending on the geography of the pair of heterospecific populations considered. We sampled a relatively low number of individuals from each site but were able to infer consistent speciation histories across distinct sample pairs, both in terms of mode of divergence and parameter estimates. This suggests that speciation histories might be reliably reconstructed using a few individuals. However, the geographical context of the study populations is also of importance, as gene flow may be heterogeneous across both species ranges. This has implications for designing and/or interpreting studies aiming at reconstructing the divergence history between two or more taxa. The present-day context of the study populations should be taken into account when interpreting parameter estimates. More specifically, species-wide interpretations should be made cautiously when, e.g., only sympatric populations are used to perform demographic history reconstruction.

## CONCLUSIONS

Here, we used genome-wide data to show that the mound-building wood ant species, *F. polyctena* and *F. aquilonia,* diverged with continuous asymmetrical gene flow across their ranges. Employing several heterospecific sample pairs and a coalescent-based inference method, we also demonstrated that the context in which present-day populations occur may influence the inference of certain demographic parameters, such as rates of migration between the populations, at different time scales. This work has important implications regarding the interpretation of studies carrying demographic history reconstruction. It will also pave the way to the study of admixture in hybrid wood ants, an emerging model in the genomics of hybridization.

## ACKNOWLEDGEMENTS

We thank CSC – IT Center for Science, Finland, for computational resources. Our work was performed under the Global Ant Genomic Alliance and was supported by an HiLIFE fellowship and an Academy of Finland grant no. 309580 to JK. BP was funded through *Societas pro Fauna et Flora Fennica* and Erasmus+ grants. VCS was funded by “Fundação Ciência e Tecnologia” (Portuguese Science Foundation grants CEECINST/00032/2018/CP1523/CT0008 and CEECIND/02391/2017). PN thanks Dominik Laetsch for bioinformatic advice and Camille Lorry for assistance with fieldwork. The authors declare no conflicts of interest.

## AUTHOR CONTRIBUTIONS

Study design: JK & PN

Data collection: AA, CB, HH, JM, JK & PN

Morphological species identification: BS

Data analysis: BP & PN

Data interpretation: BP, VCS, JK & PN

Draft writing: BP, VCS, JK & PN

Draft review and editing: all authors

Supervision: VCS, JK & PN

Project funding: JK

