## Supplementary data for "Whole-genome analysis of multiple wood ant population pairs supports similar speciation histories, but different degrees of gene flow, across their European range"

*: Co-last authors

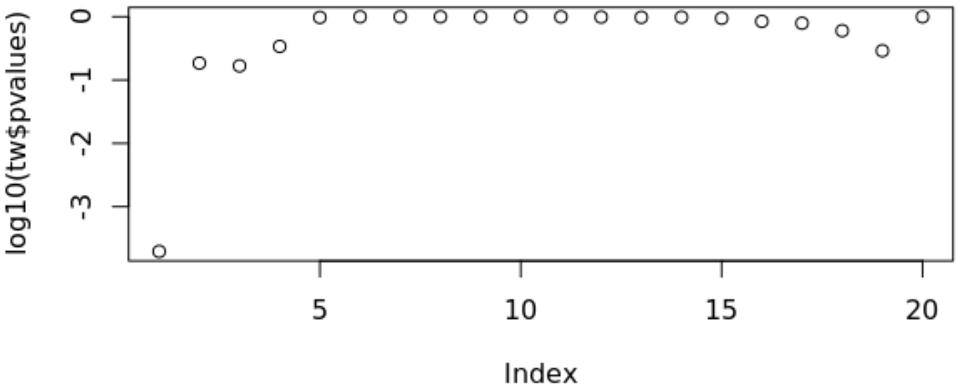


**Supplementary Figure 1 –** Tracy-Widom statistics applied to the Principal Components (PCs) assembled by the Principal Component Analysis. The first PC is determined to be statistically significant ($p$ < 0.01).

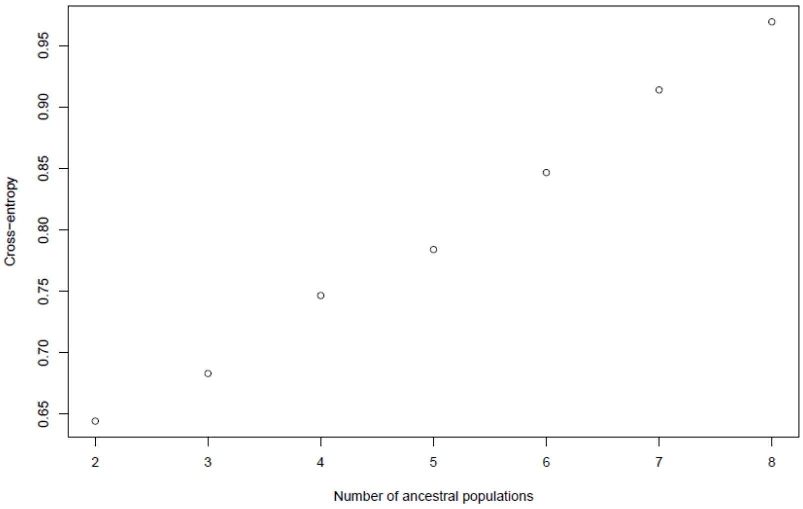


**Supplementary Figure 2 -** Cross-entropy analysis for determination of the best number of ancestral clusters (K) in the sNMF analysis. K=2 is determined to be the best number of ancestral clusters.

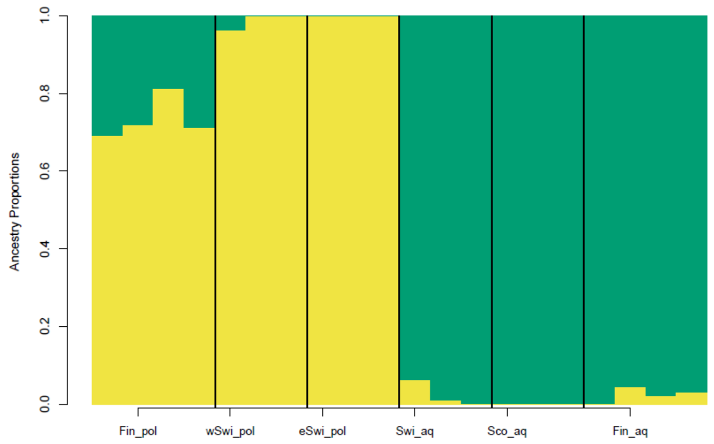


**Supplementary Figure 3 -** Ancestry proportions reconstructed by sNMF for K=2. Each bar corresponds to an individual and the different proportion of colours represent the probability of belonging to a specific cluster. Each population is separated by a black line. Abbreviations are as follows: Fin_pol = *F. polyctena* in Finland; wSwi_pol = *F. polyctena* in West Switzerland; eSwi_pol = *F. polyctena* in East Switzerland; Swi_aq = *F. aquilonia* in Switzerland; Sco_aq = *F. aquilonia* in Scotland; Fin_aq = *F. aquilonia* in Switzerland.

Note: Supplementary tables which legends are reported below can be found in the file entitled Portinha_2021_Suppl_Tables.xlsx

**Supplementary Table 1 -** Demographic parameters estimated by fastsimcoal2 in demographic model analyses. Unless bounded, the upper limit of the search range can be exceeded. Each model used only a subset of these parameters. Asterisks (*) mark parameters of models used to study the speciation history whose search ranges were altered in the "Sympatry" and "Migration after Isolation" models when testing with the Finnish comparison. The alternative minimum and maximum bounds are displayed in the appropriate columns. Double asterisks (**) mark parameters whose calculation changes between models.

**Supplementary Table 2 -** Maximum likelihood parameter estimates for all models concerning the speciation history between *Formica polyctena* and *F. aquilonia,* tested with the dataset with *F. polyctena* individuals sampled in West Switzerland and *F. aquilonia* individuals sampled in Scotland (contained 580,461 sites). All effective sizes (Ne) are given in number of haploids. Times are given in number of generations. Migration rates are scaled according to population effective sizes (2*Nm*). Maximum-likelihood estimates for parameters are taken from the run reaching the highest composite likelihood of the 100 runs performed. Likelihoods are given in logarithmic scale. Maximum observed likelihood for this dataset is -2,037,909.731. ΔLikelihood is calculated by subtracting the expected likelihood from the maximum observed likelihood.

**Supplementary Table 3 -** Maximum likelihood parameter estimates for all models concerning the speciation history between *Formica polyctena* and *F. aquilonia,* tested with the dataset with *F. polyctena* individuals sampled in East Switzerland and *F. aquilonia* individuals sampled in Scotland (contained 616,655 sites). All effective sizes (Ne) are given in number of haploids. Times are given in number of generations. Migration rates are scaled according to population effective sizes (2*Nm*). Maximum-likelihood estimates for parameters are taken from the run reaching the highest composite likelihood of the 100 runs performed. Likelihoods are given in logarithmic scale. Maximum observed likelihood for this dataset is -2,163,676.544. ΔLikelihood is calculated by subtracting the expected likelihood from the maximum observed likelihood.

**Supplementary Table 4 -** Maximum likelihood parameter estimates for all models concerning the speciation history between *Formica polyctena* and *F. aquilonia,* tested with the dataset with *F. polyctena* individuals sampled in West Switzerland and *F. aquilonia* individuals sampled in Switzerland (contained 567,578 sites). All effective sizes (Ne) are given in number of haploids. Times are given in number of generations. Migration rates are scaled according to population effective sizes (2*Nm*). Maximum-likelihood estimates for parameters are taken from the run reaching the highest composite likelihood of the 100 runs performed. Likelihoods are given in logarithmic scale. Maximum observed likelihood for this dataset is -1,993,626.486. ΔLikelihood is calculated by subtracting the expected likelihood from the maximum observed likelihood.

**Supplementary Table 5 -** Maximum likelihood parameter estimates for all models concerning the speciation history between *Formica polyctena* and *F. aquilonia,* tested with the dataset with individuals of both species sampled in Finland (contained 536,259 sites). All effective sizes (Ne) are given in number of haploids. Times are given in number of generations. Migration rates are scaled according to population effective sizes (2*Nm*). Maximum-likelihood estimates for parameters are taken from the run reaching the highest composite likelihood of the 100 runs performed. Likelihoods are given in logarithmic scale. Maximum observed likelihood for this dataset is -1,877,036.056. ΔLikelihood is calculated by subtracting the expected likelihood from the maximum observed likelihood.
